# Individuals born with congenital cataracts exhibit both persisting impairment and considerable recovery of white matter microstructure after sight restoration

**DOI:** 10.1101/2025.09.09.675092

**Authors:** Jordan D. Hassett, Cordula Hölig, Sunitha Lingareddy, Ramesh Kekunnaya, Brigitte Röder

## Abstract

**Introduction:** Individuals born with dense bilateral cataracts, for whom sight was restored later in life (congenital cataract reversal individuals), provide a unique opportunity to explore the impact of early (visual) experience on the development of the human brain.

**Methods:** Using diffusion MRI we assessed white matter microstructure in a sample of 20 congenital cataract reversal individuals using along-the-tract analysis of major visual white matter tracts and non-visual control tracts. We additionally recruited three control groups: 8 permanently congenitally blind individuals, 11 individuals with reversed later-onset cataracts (developmental cataract reversal individuals), and 24 age– and sex-matched typically sighted controls.

**Results:** Diffusion tensor metrics exhibited significant group differences which were specific to visual tracts. Compared to normally sighted controls, congenitally blind and congenital cataract reversal individuals both showed impaired white matter integrity. However, differences were much more spatially extensive for permanently congenitally blind individuals, and a direct comparison revealed relatively higher tract integrity for congenital cataract reversal individuals, suggesting a high degree of recovery following sight restoration.

**Discussion:** The present pattern of results is compatible with the idea of both a sensitive period for white matter development and significant white matter plasticity later in life. We propose that group differences are largely driven by differences in myelination. Thus, the considerable recovery in congenital cataract reversal individuals would suggest that life-long myelin plasticity may remain higher than neuronal structural plasticity.

## Introduction

While brain development is a lengthy process that continues into young adulthood, the first years of life seem to be particularly important, highlighted by remarkable neural changes that are especially relevant to function. In addition to the guiding role of genes, postnatal brain development is heavily dependent on experience^1,2^. The term ‘sensitive periods’ refers to epochs in life where brain development is especially susceptible to experience; a lack of, or aberrant experience during this phase cannot be fully reversed later in life^3^. This is distinctly evident for neural circuits in the visual system. The pioneering work of Hubel and Wiesel^4^ established in non-human animal models that a sensitive period exists for visual development. More recent studies have taken this a step further, defining multiple sensitive periods for different visual functions in non-human animals^5,6^ and humans^7–9^. The existence of such sensitive periods would suggest that early visual deprivation can considerably and permanently alter neural developmental trajectories in the visual system.

In humans, natural models for investigating the role of early visual experience on brain development are scarce. Traditionally, the impact of congenital vs. late visual deprivation on functional and structural brain organisation has been investigated in permanently blind humans^10–13^. While this research allows conclusions about sensitive periods for atypical brain development, that is, the capacity of the brain to adapt to an unexpected environment, conclusions about sensitive periods for typical brain development are not possible^13^. Thus, individuals who were born blind but recovered sight later in life must be investigated to explore whether the neural circuits can make up for the lack of early experience later in life. After undergoing cataract removal surgery, people born with dense bilateral cataracts (opaque lenses) provide such an opportunity. Clinical research in this group has consistently reported visual acuity deficits^14,15^. Cognitive neuroscience work has extended these clinical observations to higher visual functions such as coherent motion^16^ and face processing^17–19^. Despite permanent visual deficits, considerable recovery of other functions, such as biological motion and color processing, has been demonstrated as well^16,20–24^. Though limited, recent neuroimaging efforts have begun to explore the neural correlates of sight restoration after congenital visual deprivation. For instance, visual cortical morphometry (e.g. gray matter thickness and cortical surface area) was found to not recover following sight restoration^25–27^. Other studies have reported altered retinotopic organization^28^ and excitation/inhibition profiles in the visual cortex^29,30^ in individuals with reversed congenital cataracts.

Diffusion magnetic resonance imaging (MRI) can be used to compute metrics that approximate white matter microstructure: Fractional anisotropy (FA) measures the fraction of diffusion that is directional, while mean diffusivity (MD), radial diffusivity (RD) and axial diffusivity (AD), quantify the magnitude of diffusion directionally averaged, perpendicular, and parallel to fiber tracts, respectively. Despite widespread application, the neurobiological underpinnings of these diffusion metrics remain somewhat ambiguous^31^. Generally speaking, impaired white matter integrity has been associated with increased MD and RD, and decreased FA and AD. More specifically, while FA is highly sensitive to microstructural changes, it is neurobiologically nonspecific (e.g., has been related to myelin and axonal density, or in clinical cases to inflammation and edema)^31,32^. MD reflects the overall magnitude of diffusion and is nonspecific as well (e.g. changes can reflect differences in cell density or be linked to inflammation, edema or necrosis in clinical cases)^32^. In contrast, RD and AD feature much more specificity; RD is thought to be inversely linked to myelination^32,33^, while lower AD seems to reflect reduced axonal integrity^33,34^.

There is no shortage of literature surrounding DTI in permanent blindness^11^. Most studies agree that there is decreased white matter integrity in the optic radiation as a result of blindness^10,35–44^. However, the secondary findings of these studies demonstrate considerable variation. For instance, the time of onset may indeed be important for white matter microstructure, but there is some disagreement as to the exact effect. Some studies found that the optic radiation remains relatively intact for late-blind compared to early-blind individuals^35,41^, while another study instead suggested that the optic radiation may actually be more degraded in late-blind than early-blind individuals^36^. Looking at other regions, disparity also exists; where a reduction in FA in the splenium has been reported^38,40,44^, other research has found this not to be the case^42,45^. Studies have additionally found decreased white matter integrity in late visual tracts of the ventral stream like the inferior longitudinal fasciculus and inferior frontal-occipital fasciculus^10,38,42,46^, with less support for similar differences in the dorsal stream via the superior longitudinal fasciculus^10^. In summary, white matter imaging studies in permanently congenitally blind humans have consistently observed changes, including decreased FA and increased MD, in early visual tracts. Additionally, some studies have reported similar changes in higher order visual areas, including tracts of the ventral and dorsal stream (for a recent summary see Paré et al. 2023^11^).

In the only study investigating white matter integrity in congenital cataract reversal individuals^47^, the authors employed a longitudinal approach to quantify the effect of sight restoring surgery on mean FA and MD in ten early-visual, late-visual and non-visual tracts. These authors found significant surgery-related recovery in late-visual tracts, but surprisingly, they did not observe significant changes from pre-to post-surgery assessments in either the optic tract or optic radiation. In a cross-sectional analysis however, Pedersini et al. (2023)^47^ additionally found group differences between post-surgery MD values relative to a dataset retrieved from the Human Connectome Project, as the authors themselves had not scanned normally sighted control subjects.

In the present study, we used diffusion tractography to assess white matter integrity in major visual tracts for a large sample of congenital cataract reversal individuals. We opted for a more granular ‘along-the-tract’ analysis, considering that metric values are known to naturally vary along the length of a tract^31,48^. Furthermore, we additionally assessed RD and AD metrics to better unravel possible mechanisms of white matter impairments in this group. We took a cross-sectional approach involving an age– and sex-matched normally sighted control group, as well as permanently congenitally blind individuals and individuals with reversed developmental (later-onset) cataracts all recruited from the same community and scanned in the same facility. Using this approach, we hoped to isolate the effects of sight restoration after congenital visual deprivation (as opposed to permanent congenital blindness). We expected that visual tract profiles would be significantly altered in congenital cataract reversal individuals relative to normally sighted and permanently congenitally blind individuals. Specifically, we hypothesized that diffusion tensor metrics would indicate reduced white matter integrity in visual tracts for congenital cataract reversal individuals compared with normally sighted controls. However, considering the observed partial functional recovery in congenital cataract reversal individuals^21,24,49,50^, we predicted that sight restoration had induced white matter plasticity, resulting in increased integrity in congenital cataract reversal individuals compared to permanently congenitally blind individuals. Similarly, although Pedersini et al. (2023)^47^ were unable to find longitudinal changes in early-visual tracts, considering the extensive literature supporting the optic radiation as a primary tract affected by blindness^10,35–44^, we anticipated that differences between congenital cataract reversal individuals and permanently congenitally blind individuals would reflect white matter plasticity in early-visual tracts related to sight restoration.

## Methods

### Participants

Using the same original dataset as Hölig et al. (2023)^25^, additional participants were excluded (n=6) due to incomplete datasets (n=4, one congenital cataract reversal individual, three sighted controls) and issues in preprocessing (n=2, both congenitally blind individuals), leaving a final sample of 63 participants. The primary study group consisted of 20 individuals (mean age (SD) [range] = 19 (10) [6-36] years, 50% female) born with dense bilateral cataracts, surgically removed later in life (mean age at surgery (SD) [range] = 8 (9) [3 months – 31] years), henceforth referred to as congenital cataract reversal individuals (CC). The existence of congenital dense, total bilateral cataract was affirmed by a team of ophthalmologists and optometrists based on medical history, the presence of sensory nystagmus and strabismus, absence of fundus view prior to surgery and a positive family history. Full participant details are reported in supplementary table 1.

Secondary groups included eight permanently congenitally blind individuals (CB) (mean age (SD) [range] = 21 (7) [16-39] years, 37.5% female) who were totally blind (no more than rudimentary light perception) due to peripheral reasons, and 11 developmental cataract reversal individuals (DC) (mean age (SD) [range] = 16 (10) [9-43] years, 27.3% female). DC participants had a history of late onset childhood (developmental) cataracts and cataract surgery (mean age at surgery (SD) [range] = 13 (10) [7-40] years). No other sensory impairments or neurological disorders were reported by any of the participants. Full details are reported in supplementary tables 2 and 3 respectively.

A total of 24 typically sighted controls (SC) were recruited from the local community (mean age (SD) [range] = 20 (10) [6-42] years, 33.3% female). They were age– and sex-matched to CC, CB and DC participants; specifically, 16 were matched to CC individuals, eight to CB individuals, and 11 to DC individuals. Note that some SC subjects served as controls for multiple groups.

All visually impaired participants were recruited at the LV Prasad Eye Institute in Hyderabad, India. Image acquisition took place at the Lucid Medical Diagnostics radiological center in Banjara Hills in Hyderabad. Participants were given a small monetary compensation or a small gift (under age participants), and all associated costs with participating (e.g., travel, accommodation, etc.) were reimbursed. Written informed consent (orally translated into language the participant or legal guardian was able to understand) was obtained from each participant or their legal guardian prior to scanning. This study was jointly approved by the institutional ethics review board of the LV Prasad Eye Institute in Hyderabad, India, as well as the ethics commission of the German Psychological Society.

### Image Acquisition

All magnetic resonance imaging was performed using a 1.5T GE Signa HDxt scanner (GE Healthcare, Milwaukee, Wisconsin, USA) with an 8-channel head coil at the Lucid Medical Diagnostics radiological clinic in Banjara Hills, Hyderabad, India. Whole-brain diffusion weighted images were acquired with a 55-slice gradient-echo EPI2 sequence acquired in the sagittal plane, with 60 gradient directions (voxel size=0.9375mm x 0.9375mm x 2.5mm, interslice interval=2.5mm, TE=0.0936 s, TR=15.05 s, inversion time = 0, flip angle = 90 b-value = 1000 s/mm^2^, FOV = 240 x 240 x 138 mm).

### Image Processing

Initial diffusion image processing was performed using MRTrix3 (https://www.mrtrix.org/)^51^. Specifically, processing began with denoising, unringing to correct for Gibbs ringing artifacts, followed by motion and distortion correction. As reverse-phase encoded images were unavailable, synthetic ones were created using Synb0 (https://github.com/MASILab/Synb0-DISCO)^52,53^. A tensor was fitted to the preprocessed diffusion weighted image, and tensor metrics, FA, MD, RD, and AD, were then calculated for each voxel. Tensor images were registered to MNI-space, and 72 white matter tracts were segmented with TractSeg (https://github.com/MIC-DKFZ/TractSeg?tab=readme-ov-file)^54^. Of these 72 tracts, we selected a subset of 16, according to recent studies of individuals with visual impairments^10,38,46,47^. These included nine tracts related to vision; the optic radiation (OR; left and right), the splenium (corpus callosum), the inferior frontal occipital fasciculus (IFOF; left and right), the inferior longitudinal fasciculus (ILF; left and right), and the superior longitudinal fasciculus (specifically, the second division^55^(p12); SLF; left and right). Seven additional tracts were chosen as negative controls; the genu (corpus callosum), the cerebrospinal tract (CST; left and right), the cingulum (CG; left and right), and the uncinate fasciculus (UF; left and right). Tracking was performed on 10000 streamlines (a high number of streamlines reduces variability due to random seeding). We additionally used the built-in tractometry function of TractSeg^56,57^, enabling what is commonly referred to as along-the-tract statistics. Each tract was resampled at 100 points along the length of the tract, and mean FA, MD, RD, and AD values were extracted at each of those points. While many studies, including the statistical analysis of Pedersini et al. (2023)^47^, employed a ‘tract average’ approach, this might have obscured a wealth of important information; diffusion metrics vary considerably across tract profiles, as developmental or neuropathological changes will often occur in focal parts of a tract rather than uniformly across it and behavioural associations may show similar spatial specificity^48^. As such, an along-the-tract approach adds much-needed granularity and may improve sensitivity.

### Statistics

Prior to group comparisons, a modified Z-score was used to probe each group for outliers (modified z-score ≥ 3) in any of the visual tracts and metrics. For all selected tracts, group means for diffusion metrics were calculated for each of 98 segments (100 – 2 end segments). Group means were compared using independent two-sample t-tests, creating four matrices (CC vs SC participants, CB vs SC participants, DC vs SC participants, CC vs CB participants) containing the T-statistics of 16 tracts x 98 segments. Multiple comparisons correction was performed by calculating a cluster size threshold. Specifically, a maximum-cluster-size null distribution was generated for each metric (FA, MD, AD, RD) and group comparison using 10,000 permutations to calculate the maximal number of consecutive segments within a tract that demonstrated significant (p < 0.05) group differences. The maximal cluster size at the 99.9^th^ percentile of these 16 null distributions was taken as the cluster-size threshold. Observed clusters exceeding this threshold were considered significant, with a type 1 error rate of 0.001 or less.

Within each cluster, the sum of all T-statistics was taken as the cluster mass, in order to quantify the magnitude of the differences within each cluster. Additionally, in order to provide a metric of spatial specificity, the location of the cluster was recorded as the range of segments belonging to the cluster (numbered along the tract in an upstream manner (e.g. 1 is the most downstream segment)). Lastly, an average tract deviation metric was calculated for visual and non-visual tracts, by separately taking the sum of absolute values of all cluster masses for all visual tracts, and non-visual tracts respectively, then dividing by the number of tracts (nine and seven, respectively). This metric served two functions; first, summarize the total magnitude of differences between groups, and second, provide an indication of whether changes are global, or specific to visual tracts.

### Artificial intelligence usage

No generative AI tools were used at any stage of the production of this manuscript.

## Results

### Group comparisons

Significant differences were found for all four group comparisons (CC vs SC, CB vs SC, DC vs SC, CC vs CB), primarily in visual tracts (Figure 1, Table 1). Below we report the results of comparisons between CC and matched SC participants, CB and matched SC participants, and CC and CB individuals for the left and right optic radiation, the splenium, the left and right IFOF, and the left and right ILF. We additionally provide a descriptive overview of cluster location, including spatial similarity across hemispheres and comparisons. Note that DC vs SC results and discussion can be found in the supplement (supplementary figure 1, supplementary table 4).

**Figure 1:**
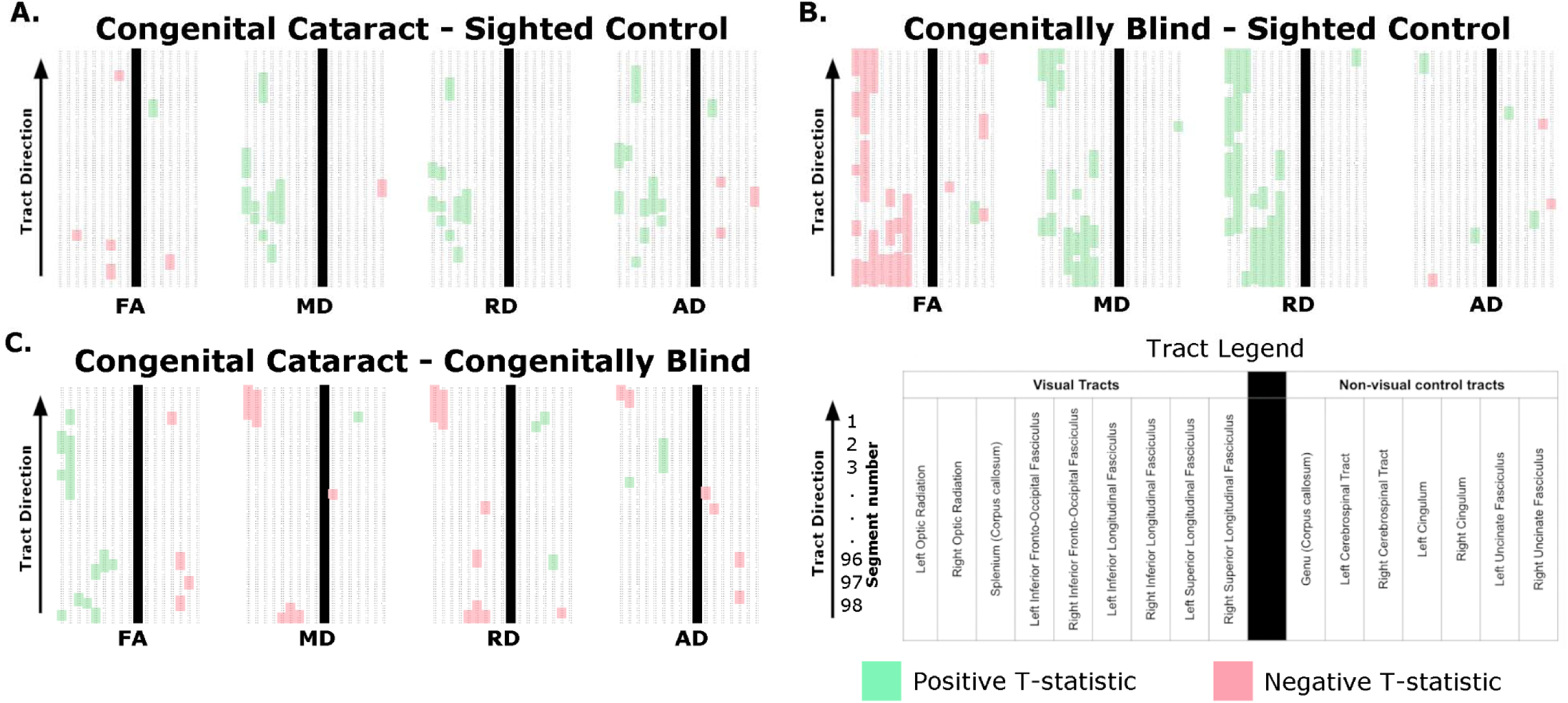
Overview of significant differences between groups. Each subplot shows significant differences in fractional anisotropy (FA), mean diffusivity (MD), radial diffusivity (RD) and axial diffusivity (AD) (labelled at the bottom). Each column represents a specific tract, for which 98 segments are displayed in rows, with downstream locations at the top. As indicated in the tract legend, tracts are organized from left to right as follows: Left Optic Radiation, Right Optic Radiation, Splenium, Left Inferior Fronto-Occipital Fasciculus, Right Inferior Fronto-Occipital Fasciculus, Left Inferior Longitudinal Fasciculus, Right Inferior Longitudinal Fasciculus, Left Superior Longitudinal Fasciculus, Right Superior Longitudinal Fasciculus, Genu, Left Corticospinal Tract, Right Corticospinal Tract, Left Cingulum, Right Cingulum, Left Uncinate Fasciculus, and Right Uncinate Fasciculus. The vertical black line indicates the separation of vision-related tracts (left) from non-visual control tracts (right). Clusters (p < 0.001) are coloured according to the direction of the difference. For example, in A, congenital cataract reversal individuals vs matched sighted controls, green clusters indicate that the respective tensor metric is significantly greater for the congenital cataract reversal group than for normally sighted controls, while red clusters indicate the opposite. This theme accordingly applies to B, permanently congenitally blind individuals vs matched sighted controls, and C, congenital cataract reversal individuals vs permanently congenitally blind individuals.

**Table 1.**
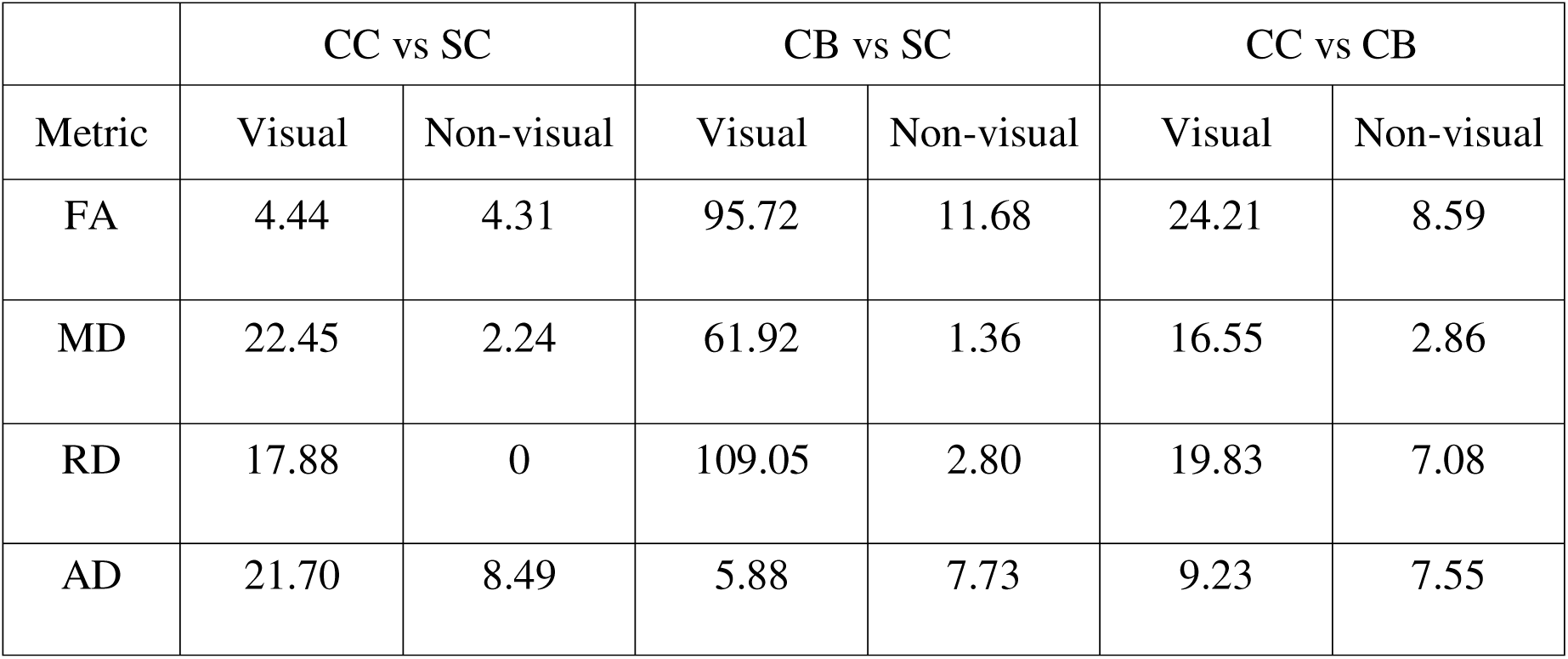
Average tract deviations of visual and non-visual tracts for each comparison and metric. **Table note:** Average tract deviation is the mean sum of cluster mass es, averaged separately across all visual, and all non-visual tracts. These scores are reported for each group comparison and tensor metric. Relatively larger values indicate that, on average, tracts showed a greater degree of differences between groups. CC – congenital cataract reversal individuals; CB – Permanently congenitally blind individuals; SC – Normally sighted control individuals; FA – Fractional anisotropy; MD – Mean diffusivity; RD – Radial diffusivity; AD – Axial diffusivity.

### Optic Radiation

Compared to SC subjects, CB individuals exhibited significantly decreased FA and increased MD, RD and AD (figure 2B, table 2). Except for FA, similar patterns of differences were found in CC compared to SC individuals (figure 2A, table 2). Comparing CC to CB participants directly revealed that FA was higher for CC individuals, while MD, RD and AD were lower (figure 2C, table 2). These differences mostly existed bilaterally, and many clusters appeared to show some degree of spatial overlap across hemispheres. Furthermore, some level of overlap was visible across comparisons, such that some significant CC vs SC and CC vs CB clusters were significant for CB vs SC too. However, there was very little apparent overlap between CC vs SC and CC vs CB comparisons.

**Figure 2:**
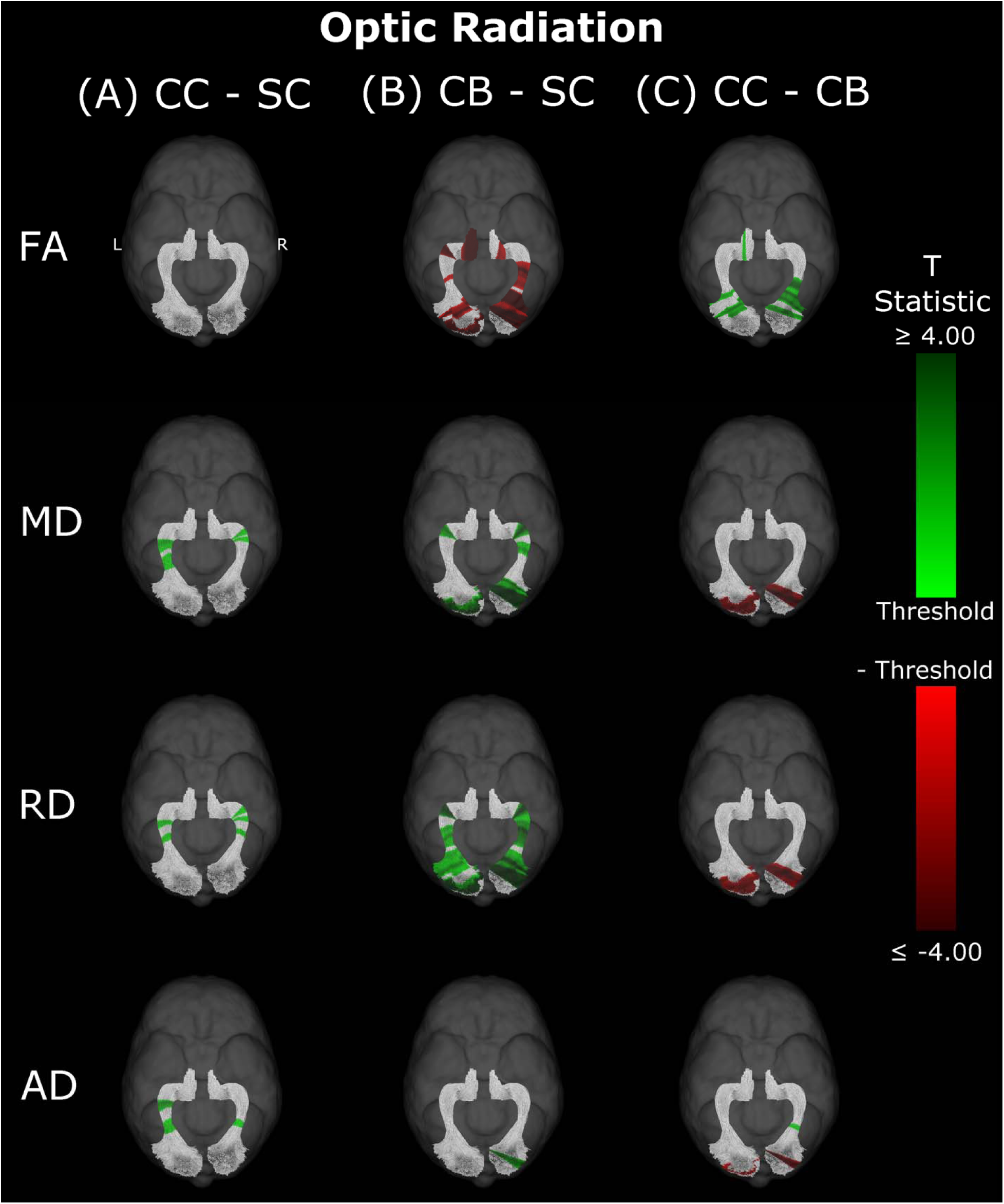
Group differences in FA, MD, RD and AD within the left and right optic radiations. The transposed segmentations of the left and right optic radiation are coloured according to the spatial location along the tract of clusters of significant differences between A) CC and SC individuals, B) CB and SC individuals, and C) CC and CB individuals. Segments coloured green indicate that the metric is higher in the first group than the second, while red segments specify the opposite. The shade of colour encodes the degree of difference, with light shades representing near-threshold T-statistics (2.032; 2.145; 2.056), and darker shades reflecting larger T-statistics. CC – congenital cataract reversal individuals; CB – Permanently congenitally blind individuals; SC – Normally sighted control individuals; FA – Fractional anisotropy; MD – Mean diffusivity; RD – Radial diffusivity; AD – Axial diffusivity.

**Table 2.** Cluster masses and segment ranges for the left and right optic radiation, by group comparison, hemisphere, and metric. **Table note:** Each entry refers to a cluster of significantly different segments found in the left or right optic radiation, organized by group comparison, hemisphere and tensor metric. Multiple entries refer to multiple clusters. Clusters are reported as ‘cluster mass [segment range]’. CC – congenital cataract reversal individuals; CB – Permanently congenitally blind individuals; SC – Normally sighted control individuals; FA – Fractional anisotropy; MD – Mean diffusivity; RD – Radial diffusivity; AD – Axial diffusivity.

| Optic Radiation |  |  |  |  |  |  |
| --- | --- | --- | --- | --- | --- | --- |
|  | CC vs SC |  | CB vs SC |  | CC vs CB |  |
| Metric | Left | Right | Left | Right | Left | Right |
| FA |  |  | 39.84 [1-11]<br>29.4 [19-28]<br>9.66 [49-52]<br>21.49 [72-77]<br>38.41 [88-98] | 142.23 [1-35]<br>76.49 [39-63]<br>24.31 [89-97] | 23.87 [20-28]<br>9.2 [36-39]<br>8.74 [94-97] | 14.83 [11-16]<br>74.83 [22-47] |
| MD | 26.11 [42-53]<br>25.68 [58-68] | 8.37 [64-67]<br>8.77 [69-72] | 43.37 [1-13]<br>20.62 [71-77] | 87.02 [1-24]<br>30.15 [51-62]<br>28.93 [69-77] | 42.36 [1-14] | 47.15 [3-17] |
| RD | 13.13 [48-53]<br>13.49 [62-67] | 10.84 [50-54]<br>8.56 [64-67]<br>8.48 [70-73] | 50.82 [1-14]<br>50.32 [21-40]<br>46.8 [44-61]<br>26.31 [71-77] | 123.95 [1-34]<br>117.95 [40-76] | 42.51 [1-15] | 45.65 [3-18] |
| AD | 22.11 [40-49]<br>23.9 [60-68] | 12.62 [41-46] |  | 22.8 [3-10] | 15.59 [1-6] | 21.71 [3-9]<br>8.78 [39-42] |

**Table 3.**
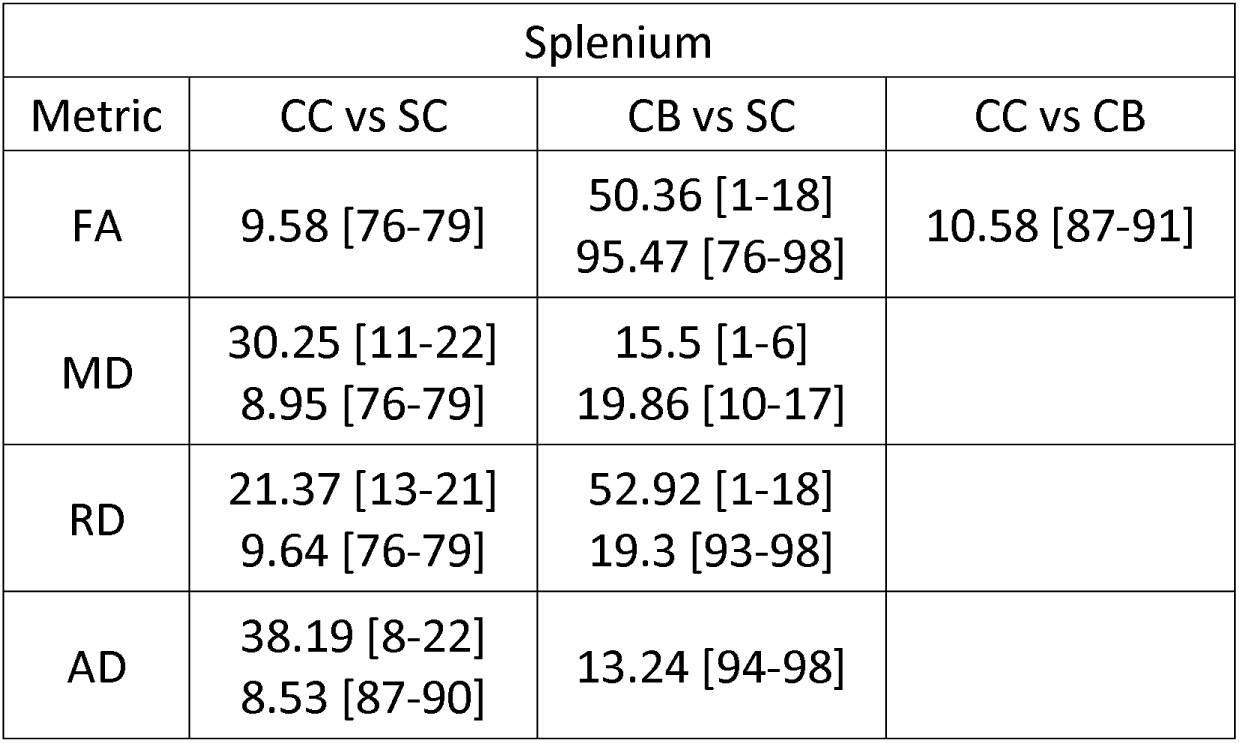
Cluster masses and segment ranges for the splenium, by group comparison and metric. **Table note:** Each entry refers to a cluster of significantly different segments found in the splenium, organized by group comparison and tensor metric. Multiple entries refer to multiple clusters. Clusters are reported as ‘cluster mass [segment range]’. CC – congenital cataract reversal individuals; CB – Permanently congenitally blind individuals; SC – Normally sighted control individuals; FA – Fractional anisotropy; MD – Mean diffusivity; RD – Radial diffusivity; AD – Axial diffusivity.

### Splenium

In CB individuals, FA and AD were significantly decreased compared to SC participants, while MD and RD were significantly increased (figure 3B, table 3). Similarly, differences in all metrics were evidenced for CC compared to SC individuals, but with AD being increased rather than decreased (figure 3A, table 3). Direct CC to CB group comparisons revealed only one small cluster of higher FA for CC individuals in the right hemispheric portion of the tract (figure 3C, table 3). Bilaterality can be partially observed, primarily in CC vs SC comparisons, with clusters in MD, RD and AD showing some degree of spatial similarity across hemispheres. Three significant clusters in CC vs SC comparisons for FA, MD and RD appeared to overlap with clusters in CB vs SC comparisons.

**Figure 3:**
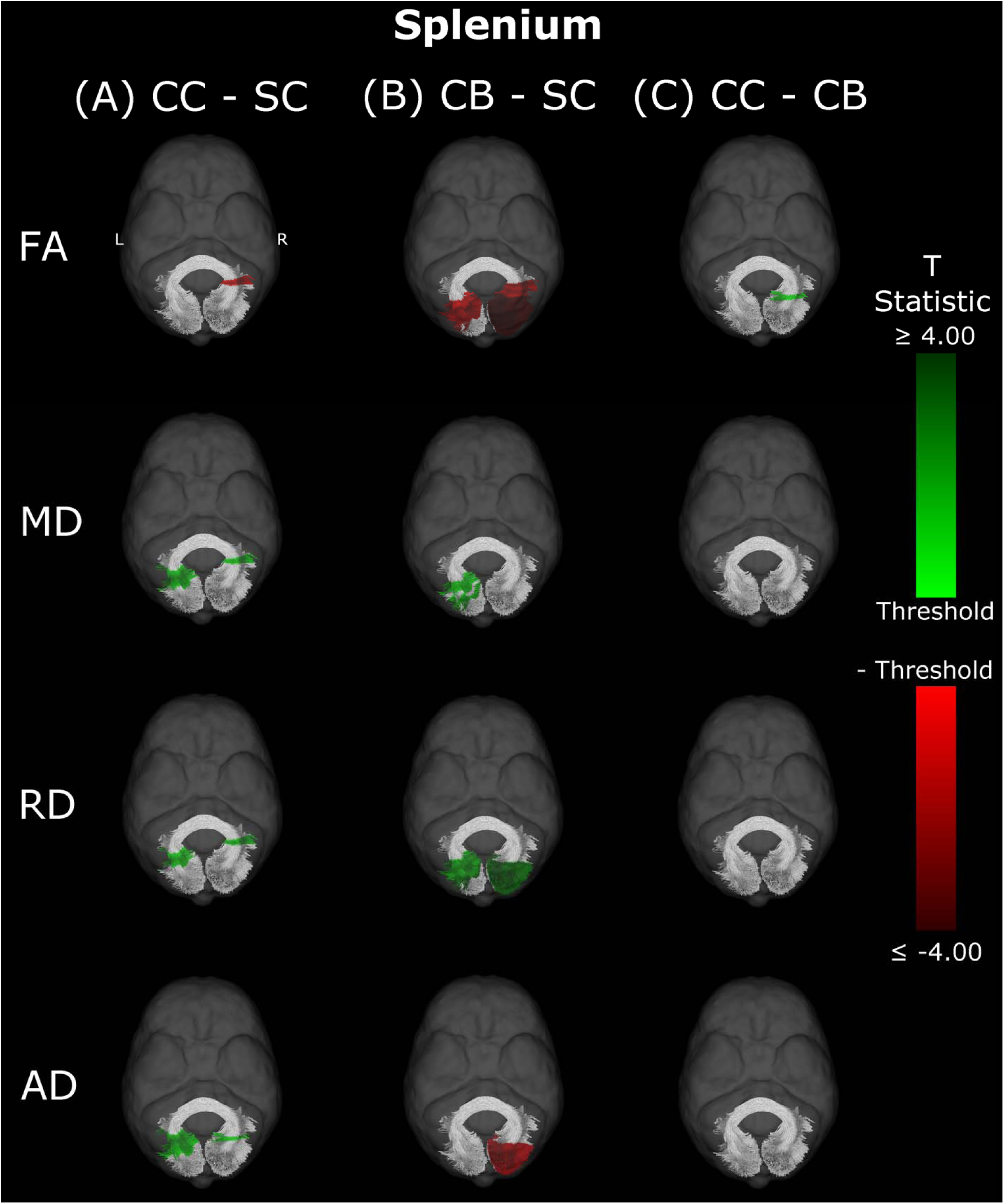
Group differences in FA, MD, RD and AD within the splenium. The transposed segmentation of the splenium is coloured according to the spatial location of clusters of significant differences between A) CC and SC individuals, B) CB and SC individuals, and C) CC and CB individuals. Segments coloured green indicate that the metric is higher in the first group than the second, while red segments specify the opposite. The shade of colour encodes the degree of difference, with light shades representing near-threshold T-statistics (2.032; 2.145; 2.056), and darker shades reflecting larger T-statistics. CC – congenital cataract reversal individuals; CB – Permanently congenitally blind individuals; SC – Normally sighted control individuals; FA – Fractional anisotropy; MD – Mean diffusivity; RD – Radial diffusivity; AD – Axial diffusivity.

### Inferior Fronto-Occipital Fasciculus

Compared with SC, CB individuals had decreased FA, and increased MD and RD (figures 4B & 5B, table 4), while CC subjects had increased MD, RD and AD (figures 4A & 5A, table 4). These differences mostly presented bilaterally and appeared to exhibit spatial similarity across hemispheres. Comparing CC and CB groups, CC individuals had significantly higher FA, lower MD and RD in the right hemisphere, but no differences in AD (figures 4C & 5C, table 4). Clusters found in CC vs SC and CC vs CB comparisons seemed to have overlap with clusters seen in CB vs SC comparisons, but not with each other.

**Figure 4:**
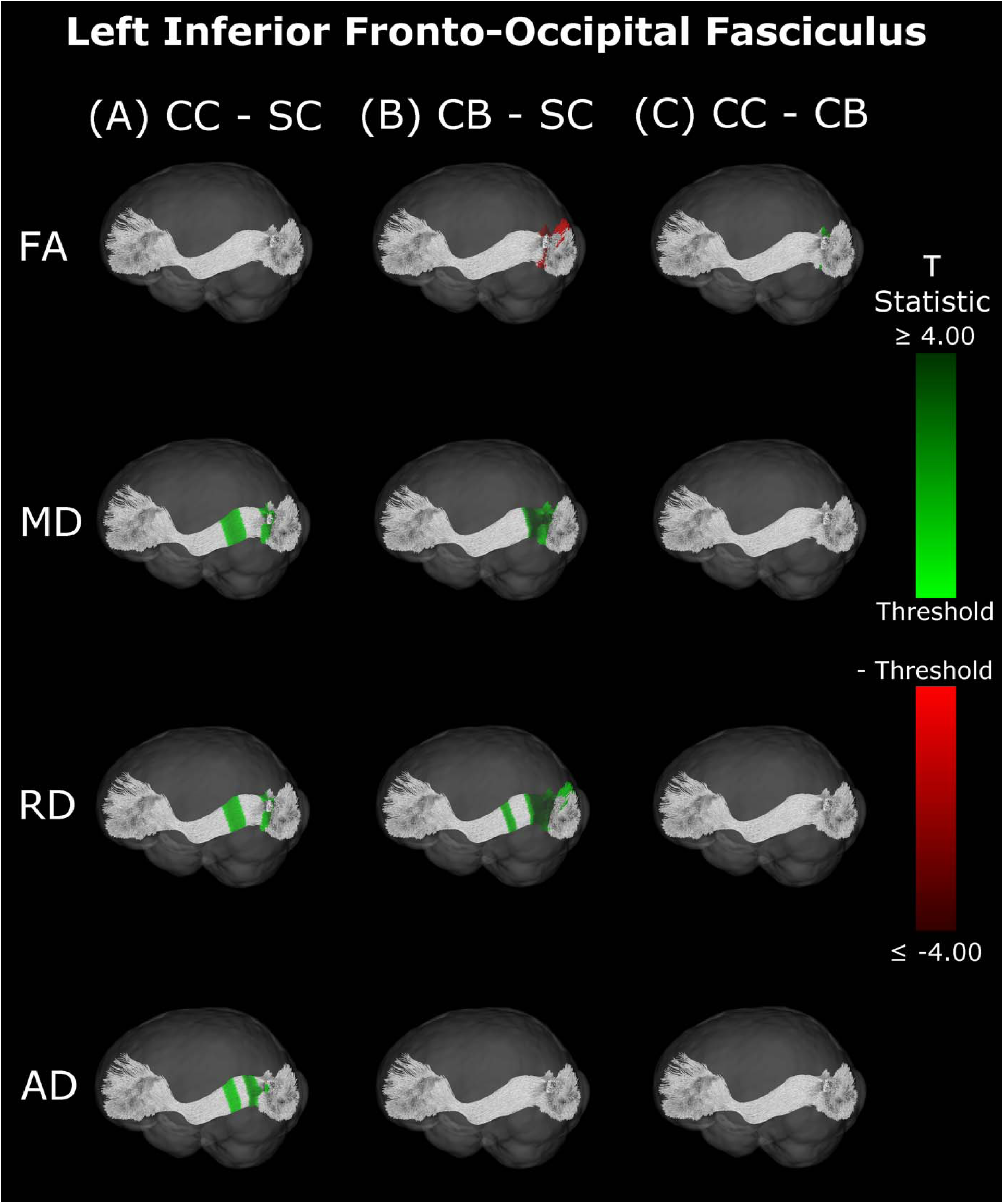
Group differences in FA, MD, RD and AD within the left inferior fronto-occipital fasciculus. The transposed segmentation of the left inferior fronto-occipital fasciculus is coloured according to the spatial location of clusters of significant differences between A) CC and SC individuals, B) CB and SC individuals, and C) CC and CB individuals. Segments coloured green indicate that the metric is higher in the first group than the second, while red segments specify the opposite. The shade of colour encodes the degree of difference, with light shades representing near-threshold T-statistics (2.032; 2.145; 2.056), and darker shades reflecting larger T-statistics. CC – congenital cataract reversal individuals; CB – Permanently congenitally blind individuals; SC – Normally sighted control individuals; FA – Fractional anisotropy; MD – Mean diffusivity; RD – Radial diffusivity; AD – Axial diffusivity.

**Figure 5:**
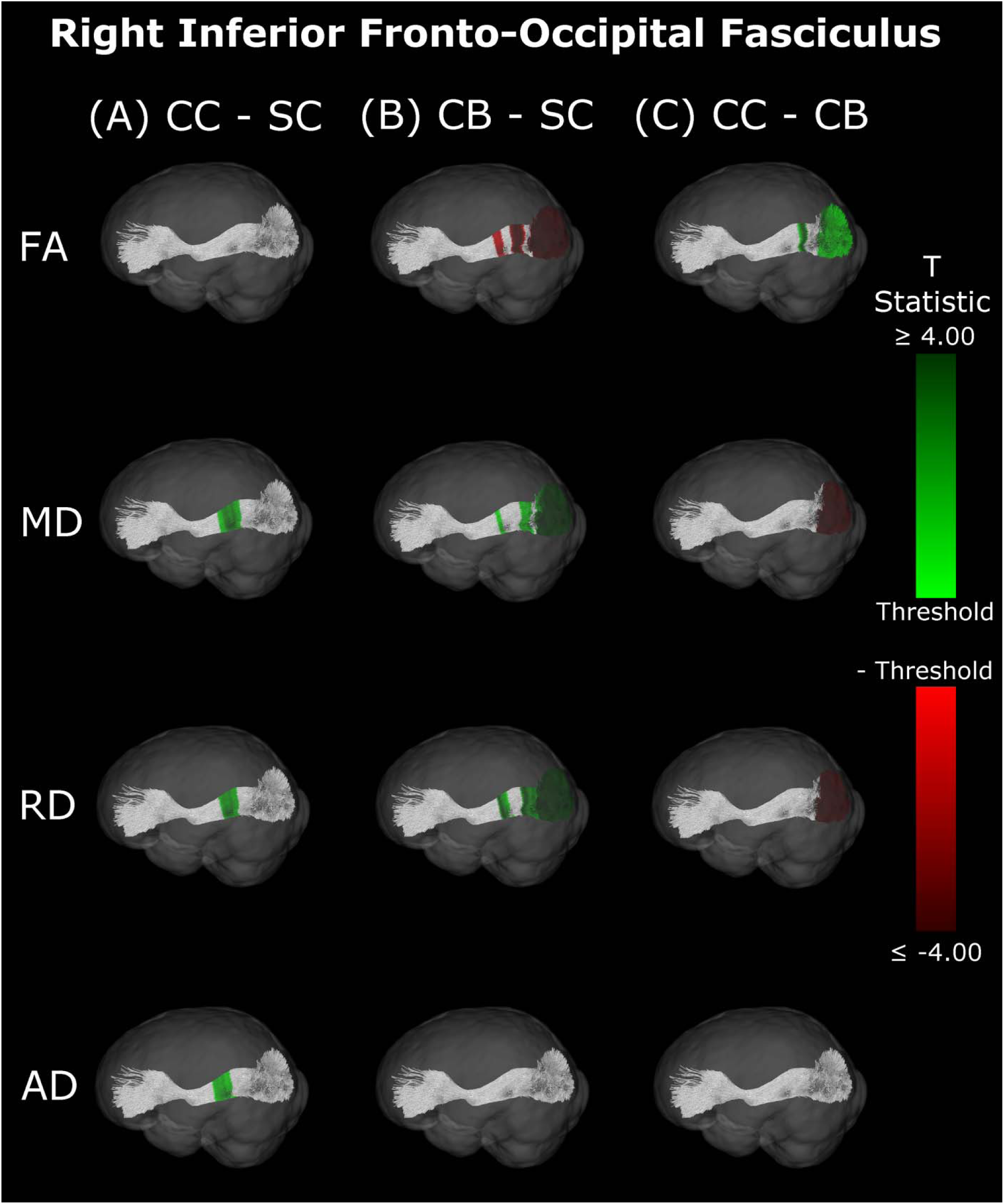
Group differences in FA, MD, RD and AD within the right inferior fronto-occipital fasciculus. The transposed segmentation of the right inferior fronto-occipital fasciculus is coloured according to the spatial location of clusters of significant differences between A) CC and SC individuals, B) CB and SC individuals, and C) CC and CB individuals. Segments coloured green indicate that the metric is higher in the first group than the second, while red segments specify the opposite. The shade of colour encodes the degree of difference, with light shades representing near-threshold T-statistics (2.032; 2.145; 2.056), and darker shades reflecting larger T-statistics. CC – congenital cataract reversal individuals; CB – Permanently congenitally blind individuals; SC – Normally sighted control individuals; FA – Fractional anisotropy; MD – Mean diffusivity; RD – Radial diffusivity; AD – Axial diffusivity.

**Table 4.** Cluster masses and segment ranges for the left and right inferior fronto-occipital fasciculi, by group comparison, hemisphere, and metric. **Table note:** Each entry refers to a cluster of significantly different segments found in the left or right optic inferior fronto-occipital fasciculus, organized by group comparison, hemisphere and tensor metric. Multiple entries refer to multiple clusters. Clusters are reported as ‘cluster mass [segment range]’. CC – congenital cataract reversal individuals; CB – Permanently congenitally blind individuals; SC – Normally sighted control individuals; FA – Fractional anisotropy; MD – Mean diffusivity; RD – Radial diffusivity; AD – Axial diffusivity.

| Inferior Fronto-Occipital Fasciculus |  |  |  |  |  |  |
| --- | --- | --- | --- | --- | --- | --- |
|  | CC vs SC |  | CB vs SC |  | CC vs CB |  |
| Metric | Left | Right | Left | Right | Left | Right |
| FA |  |  | 33.93 [86-97] | 17.86 [59-65]<br>37.76 [72-81]<br>62.53 [86-98] | 10.75 [89-92] | 18.04 [75-80]<br>16.32 [93-98] |
| MD | 29.1 [60-71]<br>15.83 [82-88] | 49 [55-72] | 57.69 [75-93] | 9.12 [59-62]<br>22.45 [77-85]<br>31.67 [89-98] |  | 15.76 [95-98] |
| RD | 23.9 [62-71]<br>13.39 [83-88] | 38.13 [59-72] | 13.16 [61-65]<br>80.84 [75-97] | 27.52 [58-66]<br>93.92 [75-98] |  | 18.72 [94-98] |
| AD | 15.38 [63-69]<br>15.38 [75-80] | 36.5 [54-68] |  |  |  |  |

### Inferior Longitudinal Fasciculus

The CB group showed decreased FA, and increased MD and RD compared to the SC group (figure 6B & 7B, table 5). Similar patterns were observed, as well as lower AD, when comparing CB to CC individuals (figure 6C & 7C, table 5). Most of these differences existed bilaterally and displayed spatial similarity across hemispheres. In contrast, CC individuals exhibited very few differences compared to SC subjects; AD in the left hemisphere was higher (figure 6A, table 5), while FA in the right hemisphere was decreased (figure 7A, table 5). The latter of these differences showed overlap with clusters in the corresponding CB vs SC comparison of FA, while nearly all clusters in CC vs CB comparisons appeared to have spatial overlap with CB vs SC clusters.

**Figure 6:**
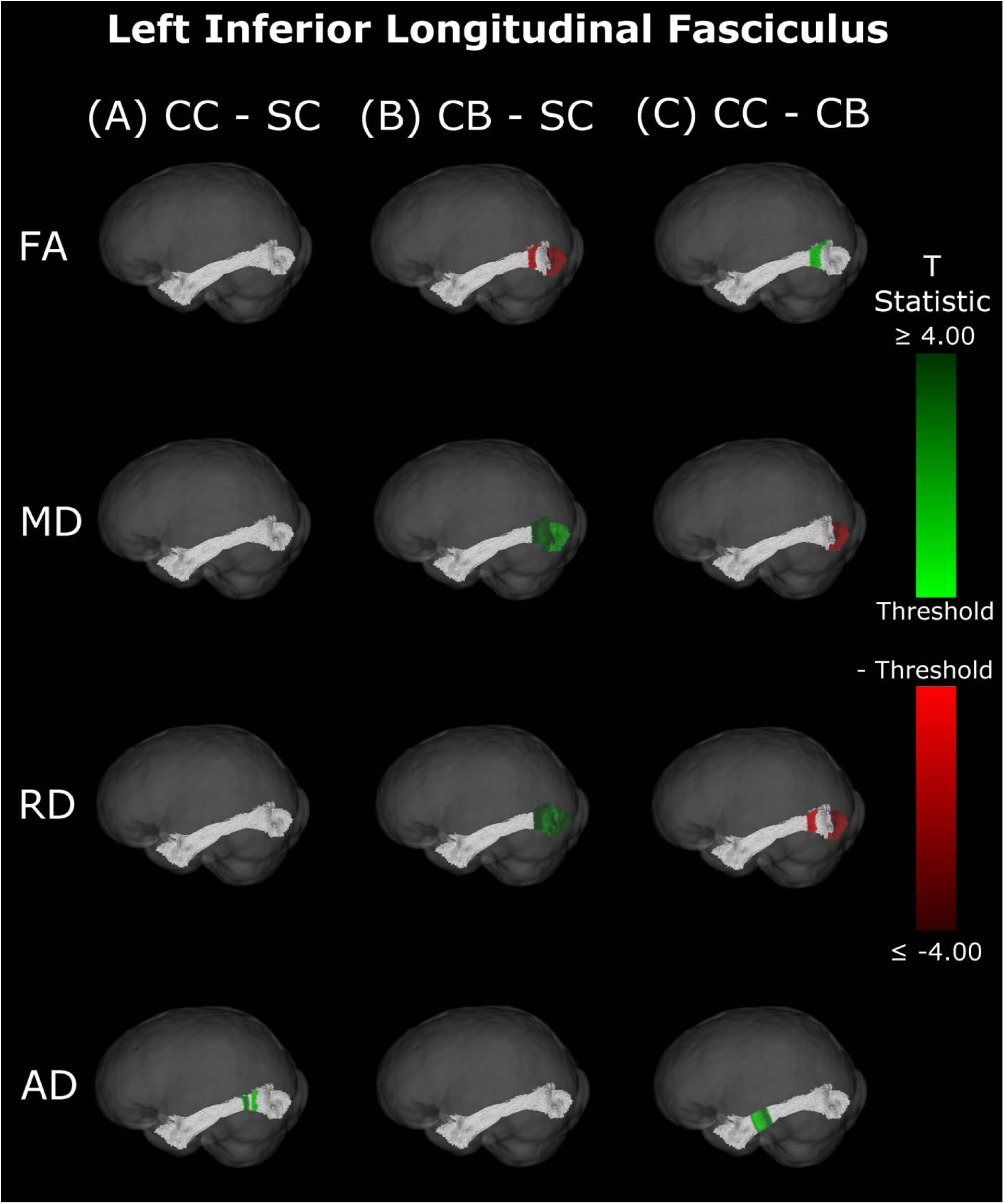
Group differences in FA, MD, RD and AD within the left inferior longitudinal fasciculus. The transposed segmentation of the left inferior longitudinal fasciculus is coloured according to the spatial location of clusters of significant differences between A) CC and SC individuals, B) CB and SC individuals, and C) CC and CB individuals. Segments coloured green indicate that the metric is higher in the first group than the second, while red segments specify the opposite. The shade of colour encodes the degree of difference, with light shades representing near-threshold T-statistics (2.032; 2.145; 2.056), and darker shades reflecting larger T-statistics. CC – congenital cataract reversal individuals; CB – Permanently congenitally blind individuals; SC – Normally sighted control individuals; FA – Fractional anisotropy; MD – Mean diffusivity; RD – Radial diffusivity; AD – Axial diffusivity.

**Figure 7:**
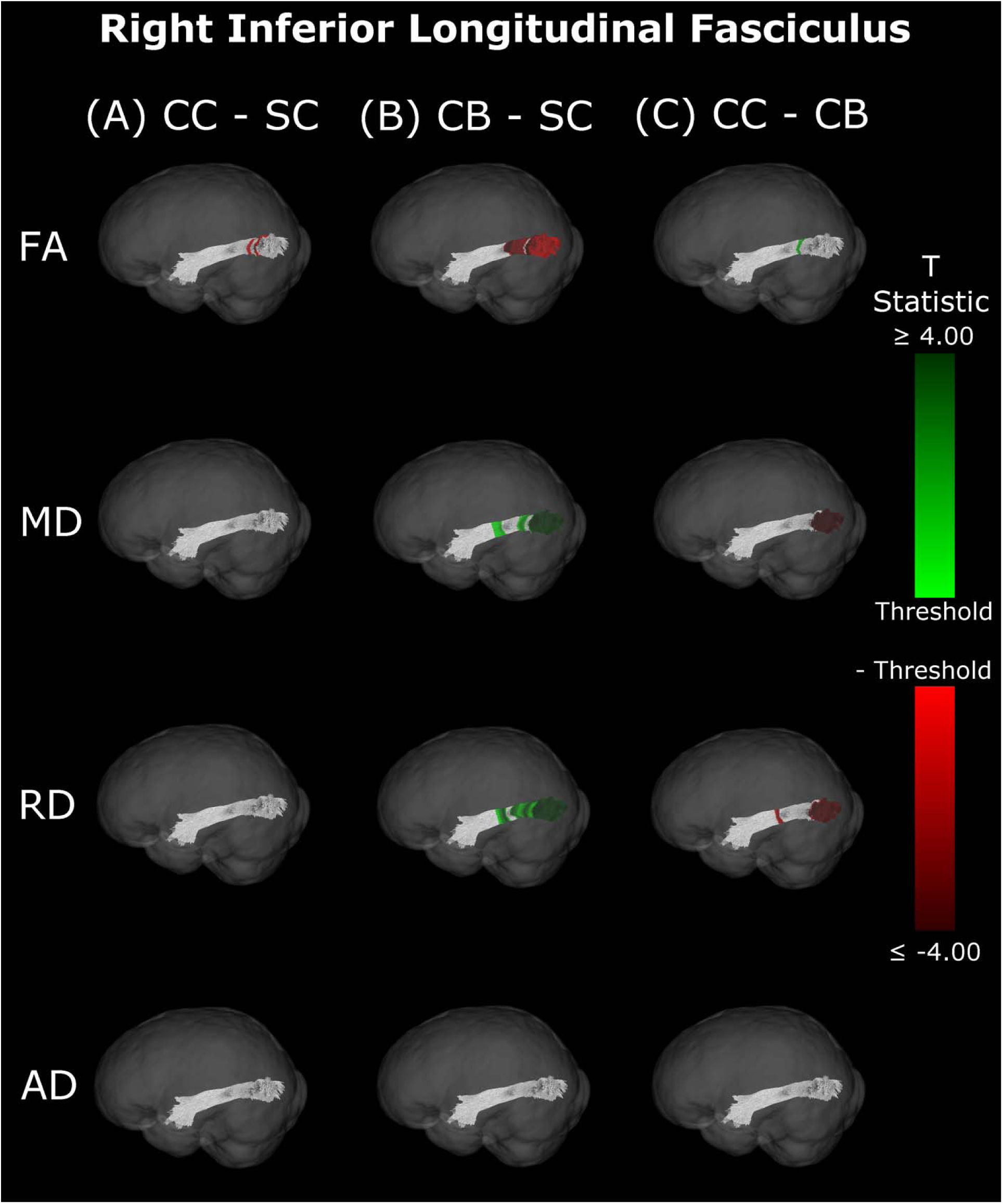
Group differences in FA, MD, RD and AD within the right inferior longitudinal fasciculus. The transposed segmentation of the right inferior longitudinal fasciculus is coloured according to the spatial location of clusters of significant differences between A) CC and SC individuals, B) CB and SC individuals, and C) CC and CB individuals. Segments coloured green indicate that the metric is higher in the first group than the second, while red segments specify the opposite. The shade of colour encodes the degree of difference, with light shades representing near-threshold T-statistics (2.032; 2.145; 2.056), and darker shades reflecting larger T-statistics. CC – congenital cataract reversal individuals; CB – Permanently congenitally blind individuals; SC – Normally sighted control individuals; FA – Fractional anisotropy; MD – Mean diffusivity; RD – Radial diffusivity; AD – Axial diffusivity.

**Table 5.**
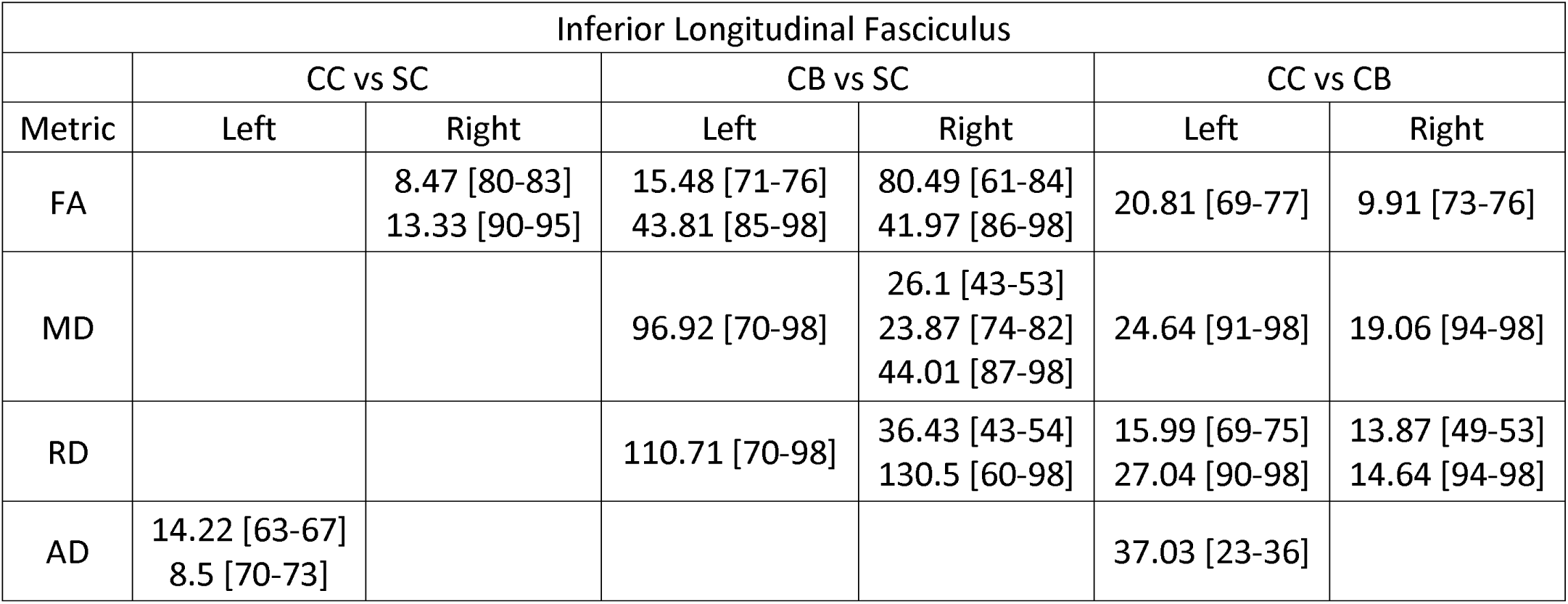
Cluster masses and segment ranges for the left and right inferior longitudinal fasciculi, by group comparison, hemisphere, and metric. **Table note:** Each entry refers to a cluster of significantly different segments found in the left or right inferior longitudinal fasciculus, organized by group comparison, hemisphere and tensor metric. Multiple entries refer to multiple clusters. Clusters are reported as ‘cluster mass [segment range]’. CC – congenital cataract reversal individuals; CB – Permanently congenitally blind individuals; SC – Normally sighted control individuals; FA – Fractional anisotropy; MD – Mean diffusivity; RD – Radial diffusivity; AD – Axial diffusivity.

| Inferior Longitudinal Fasciculus |  |  |  |  |  |  |
| --- | --- | --- | --- | --- | --- | --- |
|  | CC vs SC |  | CB vs SC |  | CC vs CB |  |
| Metric | Left | Right | Left | Right | Left | Right |
| FA |  | 8.47 [80-83]<br>13.33 [90-95] | 15.48 [71-76]<br>43.81 [85-98] | 80.49 [61-84]<br>41.97 [86-98] | 20.81 [69-77] | 9.91 [73-76] |
| MD |  |  | 96.92 [70-98] | 26.1 [43-53]<br>23.87 [74-82]<br>44.01 [87-98] | 24.64 [91-98] | 19.06 [94-98] |
| RD |  |  | 110.71 [70-98] | 36.43 [43-54]<br>130.5 [60-98] | 15.99 [69-75]<br>27.04 [90-98] | 13.87 [49-53]<br>14.64 [94-98] |
| AD | 14.22 [63-67]<br>8.5 [70-73] |  |  |  | 37.03 [23-36] |  |

### Correlations with behaviour and demographics

An exploratory analysis was performed to test whether mean metric values in the visual tracts of CC individuals were correlated with visual acuity, age at surgery, or time since surgery. For each metric, the mean of all 98 segments of each tract was calculated and was then correlated to visual acuity, age at surgery, and time since surgery, while correcting for the effect of age. Only one correlation was found to be significant (AD values in the left SLF were significantly correlated with time since surgery (p=0.037)), but this did not survive multiple comparisons correction (Spearman’s rho values are reported in supplementary tables 5, 6 and 7). Additional along-the-tract correlation analyses of the optic radiations also failed to detect significant correlations which survived statistical thresholding (not shown).

### Summary

Overall, group differences in DTI metrics were almost exclusively found in visual tracts. When these differences existed bilaterally, they displayed spatial similarity across hemispheres.

Compared to SC, CC individuals seemed to have increased MD, RD and AD in most visual tracts, but few differences in FA. Conversely, CB individuals featured extensively decreased FA relative to SC subjects, as well as increased MD and RD. Furthermore, these differences were relatively greater than those of CC vs SC participants, as evidenced by larger cluster masses.

Accordingly, a direct comparison between CC and CB individuals indicated significantly increased FA, and significantly decreased MD and RD in the CC group. Moreover, group differences between CC and SC individuals appeared to have considerable spatial overlap with the ‘large’ clusters seen in CB vs SC comparisons, particularly near the occipital terminals of tracts.

## Discussion

Congenital blindness has been demonstrated to be associated with white matter microstructure alterations in visual tracts^11^. Yet, it was unknown whether these changes are permanent or would recover if sight was restored. To test for white matter changes as a consequence of sight restoration after congenital blindness and, we acquired diffusion tensor image scans in congenitally blind individuals with restored sight due to cataract reversal surgery, permanently congenitally blind humans, individuals with reversed developmental (late-onset childhood) cataracts and age and sex matched normally sighted controls.

Results replicated previous findings in permanently congenitally blind individuals (CB group in the present study), who showed considerable degradation of white matter microstructure in visual tracts relative to SC subjects^10,11,41,42,46^. CC individuals exhibited some impairments in white matter microstructure as well, despite having underwent sight-restoring surgery. However, DTI metrics in the CC group additionally suggested a considerable degree of recovery compared to CB individuals.

### Congenital blindness and white matter impairment

Overall, DTI metrics indicated that white matter is impaired in CB relative to SC subjects. The presentation of increased MD points towards an overall increase in diffusivity within visual tracts. The concurrent finding of increased RD, with less support for widespread changes in AD, suggests that the increase in overall diffusivity is likely driven by a lower degree of myelination^32,33^. This is supported by the additionally observed decrease in FA, which indicates a decline in the ability of a tract to constrain diffusion to the primary direction. While no study to date has histologically assessed myelination in a permanently congenitally blind human population, some neuroimaging studies have speculated that there may be lower myelination in visual cortex in this population^25,58,59^.

In the present study, differences in DTI metrics were observed for the optic radiation, splenium, IFOF and ILF, but not the SLF. While the literature on permanent congenital blindness is widely varied, a large number of our findings are in line with results from previous studies. Many of these previous studies have demonstrated degradation of the optic radiation via decreased FA, increased MD or RD^10,35–39,41–44^, reinforcing the widespread bilateral changes reported in the present study. In the IFOF and ILF, support from the literature is less extensive, but still present; some studies have found decreased FA in the IFOF and ILF, but not the SLF^38,42,46^, reinforcing the current results, including the null finding in the SLF. For the splenium, some studies are consistent with the present finding of alterations^38,44^, while others did not observe differences between CB and SC individuals^42,45^.

### Persisting white matter impairment in congenital cataract reversal individuals

Importantly, while some differences seen in CC individuals relative to SC individuals mirrored those observed between CB and SC participants (although to a much lower extent), we observed qualitative differences as well. Similar to CB participants, the observed increase in MD indicates that there was an elevated level of overall diffusivity within visual tracts of CC individuals compared to the SC group. However, elevated MD was accompanied by increases in both RD and AD for the CC group, suggesting that differences were not due to a single mechanism. The absence of extensive changes in FA for CC individuals supports the idea that the overall shape of the diffusion tensor was unchanged, supported by the coinciding increase in RD and AD.

While higher RD intuitively suggests a lower degree of myelination^32,33^, AD is typically thought of as an indicator of axonal integrity^33,34^. Non-human animal work has reported that sensory deprivation induced changes in myelin were not accompanied by changes to the axons, which remained intact^60^. Although our data do not allow us to draw final conclusions about the underlying mechanisms behind higher AD in the CC group compared to the SC group, given that lower AD has previously been associated with axonal pruning^61^, we speculate that the present results may be due to reduced axonal pruning during early development or axon growth after sight restoration.

Given that only one previous study^47^ has explored white matter microstructure in congenital cataract reversal individuals, it seems crucial to compare both studies’ results where possible; Pedersini et al. (2023)^47^ explored mean FA and MD, but not RD and AD in CC individuals. In CC vs SC comparisons, our findings of increased MD and relatively unchanged FA are consistent with the cross-sectional findings of Pedersini et al. (2023)^47^. However, while we found group differences in MD to be relatively limited to the optic radiation, splenium and IFOF, these authors reported a main effect of group on MD for *all* tracts, including the non-visual tracks. This surprising finding from Pedersini et al. (2023)^47^ might be related to the absence of their own control group; these authors did not scan controls, but rather extracted control data from the Human Connectome Project – Development sample, which involved different acquisition parameters and did not include subjects from the same country (India).

### White matter recovery after sight restoration in congenital cataract reversal individuals

Compared directly to CB subjects, increased FA and decreased MD and RD seem to point towards some degree of recovery in CC individuals. Elevated FA values indicate that tracts are better able to constrain diffusion to the longitudinal axis. However, the reduction in overall diffusivity suggested by lower MD in CC individuals would indicate that group differences in FA are not purely driven by an increase in diffusion along this axis, but rather a reduction in the amount of diffusion in other axes as well. This hypothesis is supported by the concurrent observation of decreased RD. Overall, this pattern of results for the comparison of CC and CB participants would suggest that white matter recovery after sight restoration is largely driven by an increase in myelination^32,33^ of visual tracts, consistent with activity regulated mechanisms of myelin plasticity available throughout life^62^.

Remarkably, lower MD, RD and AD values were consistent across hemispheres in CC compared to CB individuals in segments where the optic radiation connects with the primary visual cortex. In contrast, no differences between the CC and SC group were observed here. These results suggest considerable recovery of thalamo-cortical connectivity in CC individuals. Pedersini et al. (2023)^47^ were unable to identify any longitudinal changes in the optic radiation, possibly due to using average values across the complete tract.

The pattern of results observed in the comparison between CC vs. CB individuals for the optic radiations was also observed for the IFOF and ILF, but not for the splenium or SLF, which is in partial agreement with the longitudinal findings of Pedersini et al. (2023)^47^. Specifically, while these authors found longitudinal increases in FA for the ILF and IFOF, they additionally observed differences in the SLF and reductions in MD of the splenium, which we did not find when comparing CC individuals to the CB group.

Our findings in the ILF are not only consistent with those of Pedersini et al. (2023)^47^, but are quite compelling in their own right. While we found large bilateral group differences for FA, MD and RD between CB and SC subjects, which was consistent with previous studies^10,38,42,46^, in CC vs SC comparisons only FA was reduced in the right ILF and only AD was increased in the left ILF. The ILF is the major white matter tract of the ventral visual stream. A recent fMRI study has identified an impressive recovery of visual categorical representations in CC individuals’ in the ventral occipito-temporal cortex compatible with the extensive white matter recovery suggested by the CC vs. CB comparison of the present study. Consistent with electrophysiological results^18,63^, Rączy et al. (2025)^64^ has recently reported largely reduced face-selectivity in CC individuals but largely unchanged selectivity for scenes. Face processing typically features a right hemispheric dominance. Thus, the lower recovery of FA in the right ILF might mirror the functional changes and observed impairments in face processing in CC individuals, while the larger pattern of ILF recovery is consistent with the high recovery of object recognition^17,19,65^.

### Spatial distribution of group differences indicates some specificity

As a particular strength of the along-the-tract approach, our results showed an impressive pattern of spatial specificity. In the comparison between CB vs SC individuals, while significant clusters can be found throughout the tract, they are very prominently located at or near the occipital terminals of all tracts. Crucially, most significant group differences between CC and CB individuals were also found near the occipital terminals, and thus overlapped with differences between CB and SC individuals. In contrast, while CC vs SC clusters did not appear in occipital areas for any tract, they did overlap with a subset of CB vs SC clusters in other locations. These results might suggest a high capacity for recovery at the occipital terminals of several visual tracts, as well as persisting impairments related to temporary congenital blindness.

### Experience-dependent myelination accounts for both impairment and recovery

Traditionally, myelin has been considered to be a rather static structure, that follows strict developmental trajectories and remains stable throughout the lifespan^66^. Prior studies have defined typical developmental trajectories for various regions of the brain, suggesting that myelination begins in the cerebellum, pons and internal capsule, then spreads to the splenium and optic radiation (3-4 months), the occipital and parietal cortices (4-6 months), and temporal and frontal cortices (6-8 months)^67^. In the optic tract, myelin can be seen (ex vivo) at term, and increases considerably in the first two years of life, then slower afterwards^68^. More recent studies have uncovered extraordinary adult myelin plasticity, providing an increasing amount of support for an activity-dependent, and thereby experience-dependent, myelination^43,66,69,70^. Neural activity is not strictly necessary for myelination to emerge in the first place during development^66^. However, studies have indicated that neuron activity can have a vast impact on myelination^71–73^. In addition to *de novo* myelination of ‘naked’ axons^74^, or the retraction of existing sheaths, myelination can additionally be remodelled through mechanisms that modulate myelin sheath length, myelin thickness, and the nodes of Ranvier^62,75–77^.

Neurons can synapse directly onto oligodendrocyte precursor cells^78^, so a lack of visual activity may reduce the production and maturation of the glial cells driving myelination in the central nervous system. A reduction in oligodendrogenesis could coincide with an increase in cortical excitability since some of the major inhibitory interneurons (e.g. parvalbumin positive GABAergic interneurons) are typically heavily myelinated^60^. A higher excitatory-inhibitory ratio in occipital cortex is a common finding in congenital blindness^29,79,80^. This ratio shows partial recovery in CC compared to CB individuals^18,29^, but future studies are needed to explore a potential link to myelin plasticity.

Structural changes following sight restoration seemingly rely on mechanisms of myelin plasticity, like *de novo* myelination or myelin remodelling^62,74–77^, via oligodendrogenesis and the proliferation of new oligodendrocyte precursor cells^81^. However, studies have suggested that differences between mechanisms of developmental myelination and later white matter plasticity exist^66,81^. Thus, we speculate that while the extensive recovery we found (compared to the CB group) might be due to mechanisms of later myelin plasticity, the persisting impairments we observed in the CC group (compared to the SC group) might indicate that adult myelin plasticity partially builds on early white matter development. While myelination can considerably increase neuronal efficiency^82^, there is a metabolic cost of development and maintenance. As such, a cost-efficient approach would not include indiscriminate myelination of all axons, but could instead partially rely on activity to determine the most important connections to myelinate, or which sheaths to maintain or improve over the complete life span.

### Limitations and future directions

Our study has a number of limitations to consider. The CB group can, at most, be an approximation of pre-surgery CC participants. While CB participants were all born blind, the causes of blindness were variable and often related to the retina. Two CB participants were additionally unable to perceive light. A longitudinal approach similar to that of Pedersini et al. (2023)^47^ would allow a more direct tracking of white matter plasticity following sight restoration. Secondly, while our sample was quite large considering the exceeding rarity of the CC population, sample size is almost always a limitation in research with rare clinical populations. Regarding the lack of correlations with variables of interest (visual acuity, age at surgery, time since surgery), we speculate that visual acuity might not be a sensitive metric for white matter changes, that persisting impairments in CC individuals may have emerged early in brain development, and that most of the recovery (observed here as null effects compared to SC individuals) occurred shortly after sight recovery. Lastly, the neurobiological interpretations of the present data rely on biomarkers that are not fully specific. While RD and AD are closely related to myelin^32,33^ and axonal integrity^33,34^ respectively, they can be influenced by other factors as well. Thus, nailing down the precise mechanisms underlying the observed white matter changes is not possible with DTI data^31,62^

## Conclusion

Ultimately, using a granular along-the-tract approach, we replicated the extensive differences between permanently congenitally blind and sighted control participants in the main visual tracts. Additionally, we found some persisting differences between normally sighted controls and congenital cataract reversal individuals, supporting the idea of a sensitive period for white matter microstructure in early visual development. However, differences between congenital cataract reversal individuals and permanently congenitally blind individuals, as well as fewer differences compared to normally sighted controls (which were highly significant for the permanently congenitally blind group) suggest that there is extensive recovery in white matter microstructure. We speculate that the main factors driving our results are related to mechanisms of myelin development and plasticity, which is consistent with the idea that life-long myelin plasticity (see as well Pedersini et al., 2023)^47^ may be higher than neuronal structural plasticity (measured via metrics of cortical morphometry^25,27^).

## Data Availability

Anonymized subject information, aggregated data and code are available at the following DOI: https://doi.org/10.25592/uhhfdm.17938. Further data can be made available upon reasonable request to the corresponding author from any qualified investigator.

## CRediT Author Contributions

**JDH:** Conceptualization, Methodology, Formal Analysis, Visualization, Writing – Original Draft. **CH:** Methodology, Data Curation, Writing – Review & Editing, Supervision. **SL:** Resources, Writing – Review & Editing. **RK:** Conceptualization, Resources, Writing – Review & Editing. **BR:** Conceptualization, Resources, Writing – Original Draft, Supervision, Project Administration, Funding Acquisition.

## Acknowledgements

We appreciate the contributions of Maria Guerreiro during study initiation, data collection and curation, and thank Seema Banerjee and Divya Jagadish for help with participant recruitment and data acquisition, as well as Armin Heinecke for an initial data quality check. We are grateful to Kabilan Pitchaimuthu, Prativa Regmi and Idris Shareef for providing clinical details of the participants during the classification process. We thank the technical staff of the Lucid Medical Diagnostics Banjara Hills in Hyderabad in India for technical assistance during collection of MRI data. We are grateful to D. Balasubramanian from the LV Prasad Eye Institute for supporting the research.

## Supplementary Information

### Participant information

**Supplementary table 1.** Congenital cataract reversal participant details (adapted from Hölig et al. (2023)^25^)

| Participant | Sex | Age (years) | Pre-surgery visual acuity <sup>a</sup> | Age at surgery (y=years, m=months) | Time since surgery (y=years, m=months) | Visual acuity (logMAR) <sup>a</sup> | Additional details |
| --- | --- | --- | --- | --- | --- | --- | --- |
| CC1 | M | 24 | N/A | 5 m | 24 y | 0.80 | Nystagmus, esotropia |
| CC2 | M | 36 | N/A | 2 y | 34 y | 0.40 | Nystagmus, esotropia |
| CC3 | M | 9 | CF at 3 m <sup>b</sup> | 7 y | 3 y | 0.80 | Nystagmus |
| CC4 | F | 6 | PL+, PR<sup>+</sup> | 4 y | 2 y | 0.90 | Nystagmus |
| CC5 | F | 18 | CFCF <sup>b</sup> | 16 y | 3 y | CF at 1 m (1.80 logMAR) | Nystagmus, exotropia, microcornea |
| CC6 | M | 32 | N/A | 6 y | 26 y | 1.30 | Nystagmus |
| CC7 | F | 28 | CFCF | 18 y | 10 y | 1.10 | Nystagmus, esotropia |
| CC8 | M | 11 | FFL | 5 m | 11 y | 0.90 | Nystagmus, esotropia |
| CC9 | M | 28 | N/A | 14 y | 15 y | 1.00 | Nystagmus, esotropia |
| CC10 | M | 13 | FFL | 1 y 3 m | 12 y | 0.30 | Nystagmus, exotropia |
| CC11 | M | 32 | N/A | 29 y 5 m | 3 y | 1.20<sup>e</sup> | Nystagmus, esotropia |
| CC12 | F | 34 | 1.52 <sup>c</sup> | 31 y 4 m | 3 y 4 m | 1.1 | Nystagmus, exotropia |
| CC13 | F | 26 | N/A | 8 m | 26 y | 0.80 | Exotropia <sup>d</sup> |
| CC14 | F | 10 | FFL | 3 y 6 m | 7 y | 0.40 | Nystagmus, exotropia |
| CC15 | F | 7 | FFL | 3 m | 7 y | 0.80 | Nystagmus, esotropia |
| CC16 | F | 17 | CFCF <sup>b</sup> | 10 y 3 m | 7 y | 1.48<sup>a</sup> | Nystagmus |
| CC17 | F | 11 | CF at 1 m <sup>e</sup> | 10 y 7 m | 6 m | 1.30 | Nystagmus |
| CC18 | M | 8 | FFL <sup>b</sup> | 5 y 4 m | 3 y | 0.40 | Nystagmus, esotropia |
| CC19 | F | 18 | FFL at 1 m | 2 y 1 m | 16 y | 0.48 | Nystagmus, esotropia, microcornea |
| CC20 | M | 13 | CF at 0.5 m | 5 y 4 m | 8 y | 0.74 | Nystagmus, esotropia |

Note. M□=□male; F□=□female; N/A□=□not available; FFL□=□fixing and following light; PL+□=□able to perceive light; PR+□=□able to report the location of light; CF□=□counting fingers; CFCF□=□counting fingers close to face. <sup>a</sup>Positive logMAR values indicate the degree of vision loss, i.e., higher values denote lower visual acuity. <sup>b</sup>(Partially) absorbed cataracts. <sup>c</sup>Absorbed cataracts. <sup>d</sup>The presence of nystagmus was not reported in the medical file. <sup>e</sup>Neither retinal glow with retinoscopy nor fundus view with ophthalmoscopy was obtained prior to cataract surgery.

**Supplementary table 2.** Congenitally blind participant details (adapted from Hölig et al. (2023)^25^)

| Participant | Sex | Age (years) | Visual acuity | Cause of blindness |
| --- | --- | --- | --- | --- |
| CB1 | M | 39 | N/A | Microphthalmia OU |
| CB2 | M | 21 | NLP | Microphthalmia OU, microcornea OU |
| CB3 | F | 19 | PL<sup>+</sup> | Microphthalmia OU |
| CB4 | M | 17 | PL<sup>+</sup> | Phthisis bulbi OD, anterior staphyloma OS |
| CB5 | F | 19 | PL<sup>+</sup> | Anterior staphyloma OD, phthisis bulbi OS |
| CB6 | M | 23 | CFCF | Corneal scar OU, microphthalmia OU |
| CB7 | M | 16 | PL<sup>+</sup>, PR<sup>+</sup> | Microphthalmia OU, band-shaped Keratopathy OU |
| CB8 | F | 17 | NLP | Leber's congenital amaurosis |

Note. M□=□male; F□=□female; OU□=□both eyes; OD□=□right eye; OS□=□left eye; N/A□=□not available; FFL□=□fixing and following light; NLP□=□no light perception; PL+□=□able to perceive light; PR+□=□able to report the location of light; CFCF□=□counting fingers close to face.

**Supplementary table 3.** Developmental cataract participant details (adapted from Hölig et al. (2023)^25^)

| Participant | Sex | Age (years) | Cataract onset <sup>a</sup> | Pre-surgery visual acuity <sup>b</sup> | Age at surgery (y = years, m = months) | Time since surgery (y = years, m = months) | Visual acuity <sup>b</sup> |
| --- | --- | --- | --- | --- | --- | --- | --- |
| DC1 | M | 13 | Developmental | PL+, PR+ | 9 y | 4 y | 1.78 |
| DC2 | M | 13 | Developmental | 0.80 | 12 y 6 m | 7 m | 0.10 |
| DC3 | F | 43 | Age-related <sup>c</sup> | 0.60 | 40 y 4 m <sup>d</sup> | 3 y | 0.30 |
| DC4 | M | 12 | Congenital | 0.80 | 11 y 10 m | 8 m | 0.60 |
| DC5 | M | 9 | Developmental | 1.04 | 7 y 2 m | 2 y | 0.30 |
| DC6 | M | 9 | Developmental | 0.90 | 7 y | 2 y | 0.10 |
| DC7 | M | 17 | Developmental | 0.40 | 12 y 6 m | 5 y | 0.10 |
| DC8 | M | 13 | Developmental | 1.00 | 8 y | 5 y | 0.20 |
| DC9 | M | 11 | Developmental | 0.48 | 8 y 4 m | 4 y | 0.30 |
| DC10 | F | 13 | Developmental | 0.50 | 10 y 2 m | 3 y | 0.10 |
| DC11 | F | 21 | Developmental | 0.40 | 17 y 4 m | 4 y | 0.20 |

*Note.* M = male; F = female; PL+ = able to perceive light; PR+ = able to report the location of light, - = neither nystagmus nor strabismus reported in the medical records. <sup>a</sup> Developmental cataracts typically develop gradually and remain unnoticed for some time; thus, it is not possible to define a clear onset. <sup>b</sup> Positive logMAR values indicate the degree of vision loss, i.e., higher values denote lower visual acuity. <sup>c</sup> The onset of cataracts was after the age of 12 years. <sup>d</sup> Operated only in one eye.

## Results

### DC vs SC comparisons

DC individuals showed higher MD, RD and AD compared to SC individuals (supplementary figure 1C, supplementary table 4). As with other group comparisons, clusters were primarily seen in visual tracts, which again exhibited higher average tract deviation (MD: 20.39, RD: 16.19, AD: 20.85) than non-visual tracts (MD: 1.80, RD: 2.02, AD: 8.67). More precisely, differences were primarily observed for early visual tracts, namely the left and right OR and the splenium, although the left IFOF also showed differences for all four metrics. As with CC individuals, average tract deviation scores indicated relatively fewer differences between DC and SC individuals for FA (visual: 8.38, non-visual: 1.82), but differences in the left and right ILF may be noteworthy.

**Supplementary table 4.**
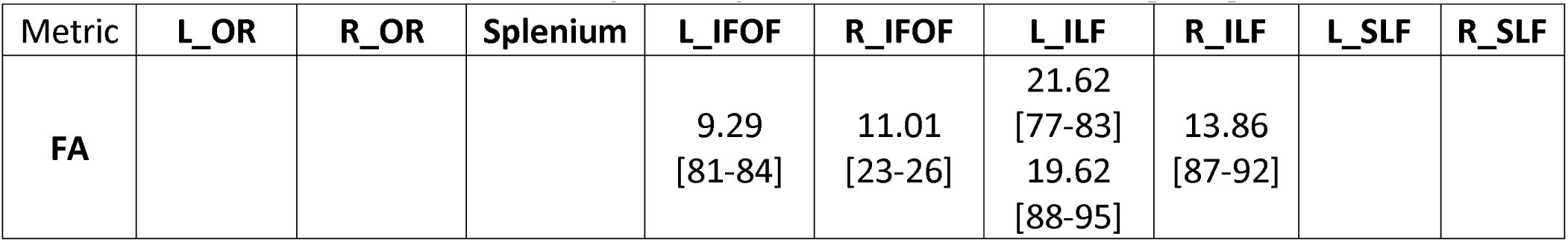
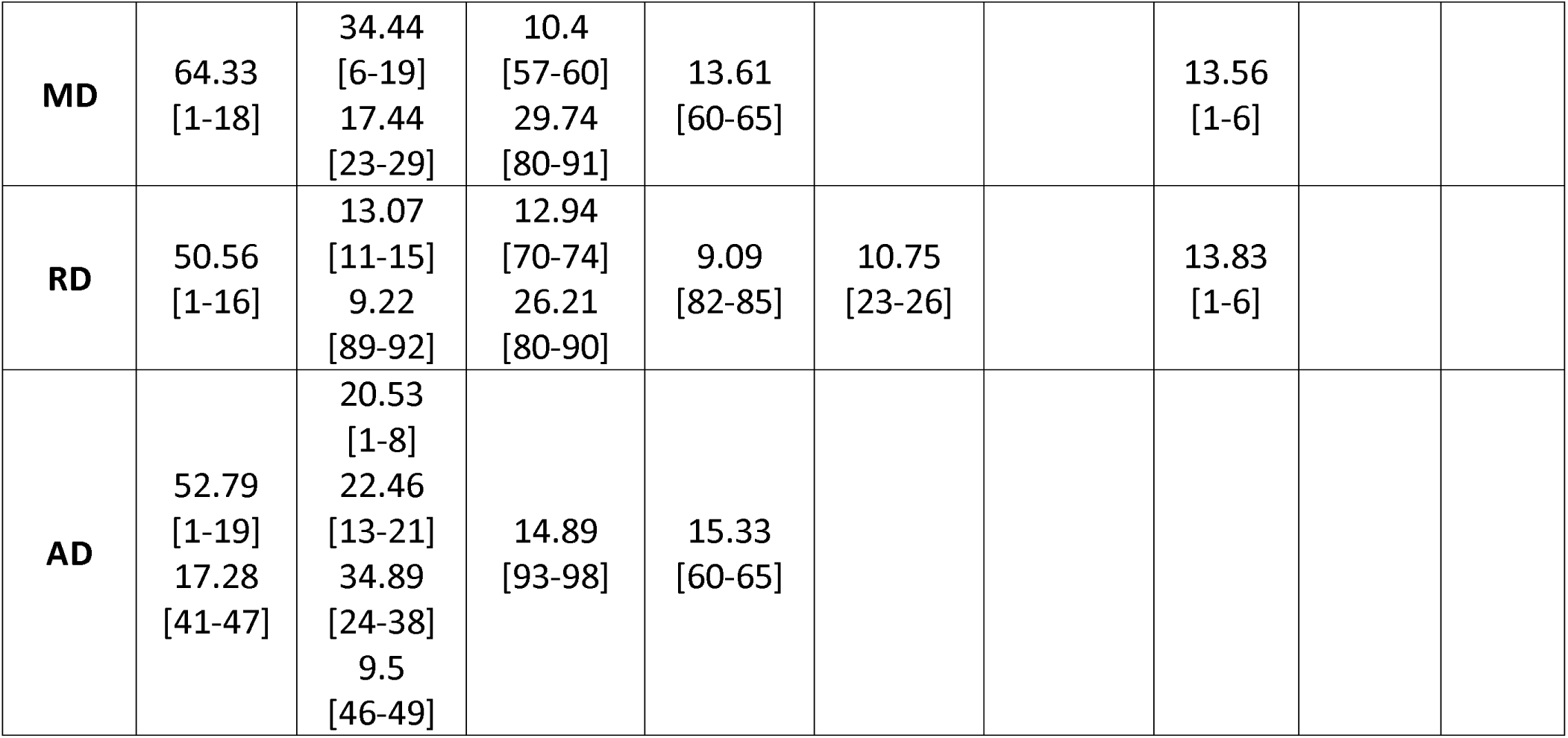
Cluster masses and segment ranges for visual tracts of DC vs SC participants. **Table note**: Each entry refers to a cluster of significantly different segments found in the left or right inferior longitudinal fasciculus, organized by group comparison, hemisphere and tensor metric. Multiple entries refer to multiple clusters. Clusters are reported as ‘cluster mass [segment range]’. CC – congenital cataract reversal individuals; CB – Permanently congenitally blind individuals; SC – Normally sighted control individuals; FA – Fractional anisotropy; MD – Mean diffusivity; RD – Radial diffusivity; AD – Axial diffusivity.

**Supplementary figure 1:**
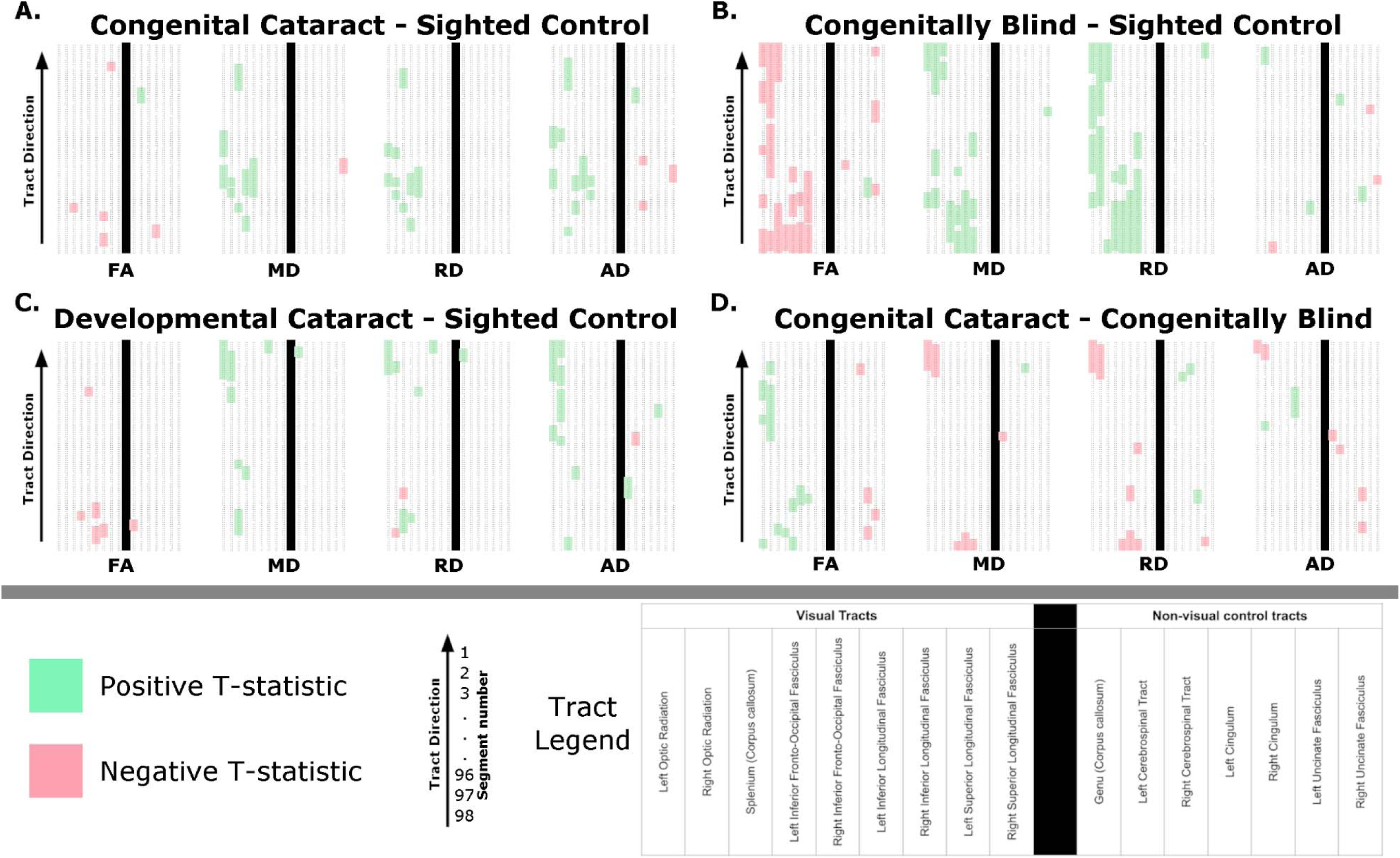
Overview of significant differences between groups. Each subplot shows significant differences in fractional anisotropy (FA), mean diffusivity (MD), radial diffusivity (RD) and axial diffusivity (AD) (labelled at the bottom). Each column represents a specific tract, for which 98 segments are displayed in rows, with downstream locations at the top. As indicated in the tract legend, tracts are organized from left to right as follows: Left Optic Radiation, Right Optic Radiation, Splenium, Left Inferior Fronto-Occipital Fasciculus, Right Inferior Fronto-Occipital Fasciculus, Left Inferior Longitudinal Fasciculus, Right Inferior Longitudinal Fasciculus, Left Superior Longitudinal Fasciculus, Right Superior Longitudinal Fasciculus, Genu, Left Corticospinal Tract, Right Corticospinal Tract, Left Cingulum, Right Cingulum, Left Uncinate Fasciculus, and Right Uncinate Fasciculus. The vertical black line indicates the separation of vision-related tracts (left) from non-visual control tracts (right). Clusters (p < 0.001) are coloured according to the direction of the difference. For example, in A, congenital cataract reversal individuals vs matched sighted controls, green clusters indicate that the respective tensor metric is significantly greater for the congenital cataract reversal group than for normally sighted controls, while red clusters indicate the opposite. This theme accordingly applies to B, permanently congenitally blind individuals vs matched sighted controls, C, developmental cataract reversal individuals vs matched normally sighted controls, and D, congenital cataract reversal individuals vs permanently congenitally blind individuals.

### Correlations with variables of interest

**Supplementary table 5:**
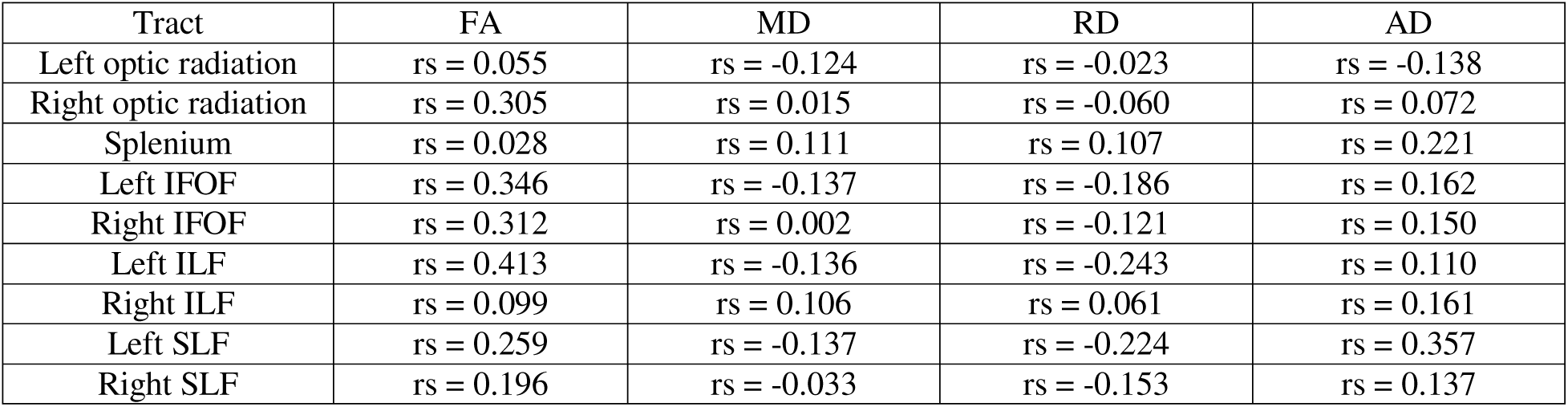
Spearman’s rho values for the relationship between age at surgery and mean DTI metrics of visual tracts. **Table note:** Spearman’s rho values are reported for the correlation between each tract’s mean metric values and age at surgery in months. Only congenital cataract reversal individuals were included. No significant correlations were revealed. FA – Fractional anisotropy; MD – Mean diffusivity; RD – Radial diffusivity; AD – Axial diffusivity; IFOF – inferior fronto-occipital fasciculus; ILF – inferior longitudinal fasciculus; SLF – superior longitudinal fasciculus.

**Supplementary table 6:**
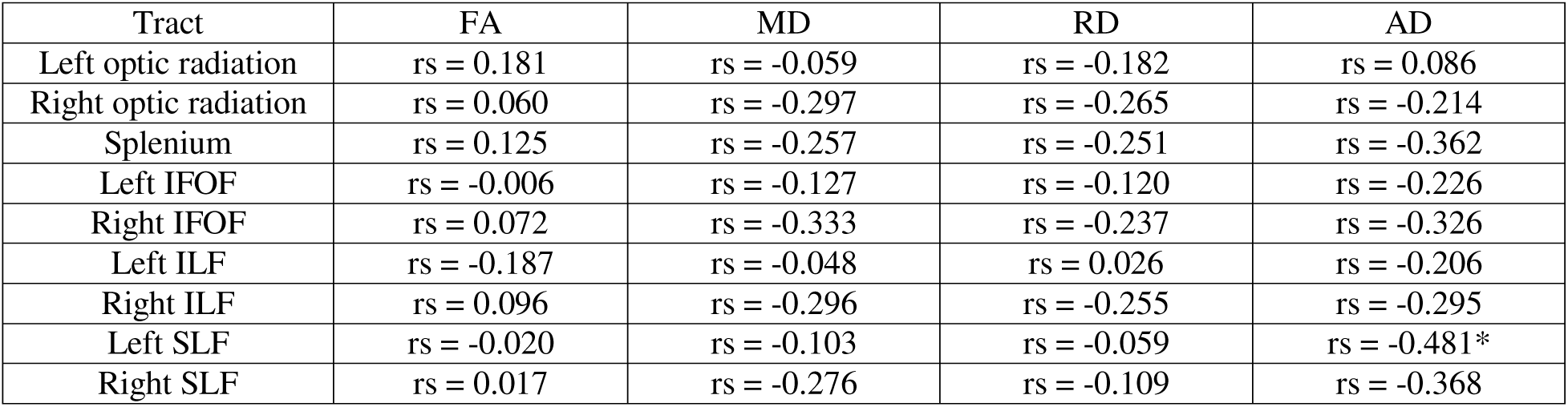
Spearman’s rho values for the relationship between time since surgery and mean DTI metrics of visual tracts. **Table note:** Spearman’s rho values are reported for the correlation between each tract’s mean metric values and time since surgery in months. Only congenital cataract reversal individuals were included. AD values in the left SLF were significantly correlated to time since surgery (p = 0.037), but this did not survive multiple comparisons correction. FA – Fractional anisotropy; MD – Mean diffusivity; RD – Radial diffusivity; AD – Axial diffusivity; IFOF – inferior fronto-occipital fasciculus; ILF – inferior longitudinal fasciculus; SLF – superior longitudinal fasciculus.

**Supplementary table 7:** Spearman’s rho values for the relationship between visual acuity and mean DTI metrics of visual tracts. **Table note:** Spearman’s rho values are reported for the correlation between each tract’s mean metric values and visual acuity reported as logMAR values (higher values indicate lower visual acuity). Only congenital cataract reversal individuals were included. No significant correlations were revealed. FA – Fractional anisotropy; MD – Mean diffusivity; RD – Radial diffusivity; AD – Axial diffusivity; IFOF – inferior fronto-occipital fasciculus; ILF – inferior longitudinal fasciculus; SLF – superior longitudinal fasciculus.

| Tract | FA | MD | RD | AD |
| --- | --- | --- | --- | --- |
| Left optic radiation | rs = -0.023 | rs = -0.001 | rs = 0.105 | rs = -0.165 |
| Right optic radiation | rs = 0.180 | rs = 0.129 | rs = 0.069 | rs = 0.052 |
| Splenium | rs = 0.373 | rs = -0.112 | rs = -0.201 | rs = 0.006 |
| Left IFOF | rs = 0.233 | rs = 0.004 | rs = -0.025 | rs = 0.176 |
| Right IFOF | rs = 0.226 | rs = 0.057 | rs = -0.013 | rs = 0.050 |
| Left ILF | rs = 0.265 | rs = 0.031 | rs = -0.085 | rs = 0.085 |
| Right ILF | rs = 0.101 | rs = 0.065 | rs = 0.040 | rs = 0.190 |
| Left SLF | rs = -0.075 | rs = -0.081 | rs = -0.057 | rs = 0.086 |
| Right SLF | rs = 0.058 | rs = 0.024 | rs = -0.041 | rs = 0.104 |

## Discussion

### Alterations to white matter in developmental cataract reversal individuals

We consider the DC group to be a control group for the treatment (cataract surgery) rather than an experimental group, e.g., to determine the timing of sensitive periods. The main reasons are: First and foremost, it is important to note that the DC group is extremely variable. Inherently, developmental cataracts happen gradually and can go unnoticed for some time. Thus, defining a time of onset is impossible. Second, as can be seen in supplementary table 3, the majority of participants had some level of vision prior to surgery, as intervention is typically sought prior to complete loss of vision.

Relative to matched sighted controls, we observed increases in MD, RD and AD mostly in the optic radiation and a few differences in FA in the left and right ILF. Changes in the optical radiation were prominent at the occipital terminals of the optic radiation, whereas in CC participants, alterations compared to the SC group were more central in tracts. While we refrain from interpreting the results for the optic radiation of the DC group, we noticed that they qualitatively differed from those of the CC group (location and parameter). Interestingly are the overlaps of FA decreases in the ILF, which seem even more extensive in the DC group. This result could indicate a protracted vulnerability of the ventral occipital cortex in childhood. The functional consequences are yet unknown. The only existing study on face processing found reliable impairments for CC individuals but not for DC individuals^18^.

